# Automated and customizable quantitative image analysis of whole *C. elegans* germlines

**DOI:** 10.1101/2020.11.25.398685

**Authors:** Erik Toraason, Victoria L. Adler, Nicole A. Kurhanewicz, Acadia DiNardo, Adam M. Saunders, Cori K. Cahoon, Diana E. Libuda

## Abstract

Arranged in a spatial-temporal gradient for germ cell development, the adult germline of *Caenorhabditis elegans* is an excellent system for understanding the generation, differentiation, function, and maintenance of germ cells. Imaging whole *C. elegans* germlines along the distal-proximal axis enables powerful cytological analyses of germ cell nuclei as they progress from the pre-meiotic tip through all the stages of meiotic prophase I. To enable high-throughput image analysis of whole *C. elegans* gonads, we developed a custom algorithm and pipelines to function with image processing software that enables: 1) quantification of cytological features at single nucleus resolution from immunofluorescence images; and, 2) assessment of these individual nuclei based on their position within the germline. We demonstrate the capability of our quantitative image analysis approach by analyzing multiple cytological features of meiotic nuclei in whole *C. elegans* germlines. First, we quantify double strand DNA breaks (DSBs) per nucleus by analyzing DNA-associated foci of the recombinase RAD-51 at the single-nucleus resolution in the context of whole germline progression. Second, we quantify the DSBs that are licensed for crossover repair by analyzing foci of MSH-5 and COSA-1 when they associate with the synaptonemal complex during meiotic prophase progression. Finally, we quantify P-granule composition across the whole germline by analyzing the colocalization of PGL-1 and ZNFX-1 foci. Our image analysis pipeline is an adaptable and useful method for researchers spanning multiple fields utilizing the *C. elegans* germline as a model system.

## Introduction

Reproduction in many sexually reproducing organisms requires the formation of haploid gametes. Gametes originate from germ cells that divide and differentiate to generate a germline, which is also known as the “totipotent” or “immortal” cell lineage due to its ability to pass on its genetic information to the next generation [1]. Studies of germ cells in multiple systems have revealed molecular mechanisms of germ cell development, function, and maintenance. Over the past several decades, the use of genetics and cytology has been instrumental for understanding fundamental aspects of germ cell biology.

For germ cell studies, the *Caenorhabditis elegans* germline provides unique manipulation and visualization advantages [2,3]. In adult hermaphrodites, there are two complete tube-shaped gonads each form a U-shape when contained within the adult animal [1]. Within the adult hermaphrodite germline, ~1000 germ cell nuclei are positioned around the circumference of the tube and are arranged in a spatial-temporal gradient according to developmental stage along the distal-proximal axis. At the distal end of the gonad (pre-meiotic tip or proliferative zone), mitotically-cycling nuclei move proximally until they reach the leptotene/zygotene region that commits them to enter meiosis, the specialized cell division that generates haploid gametes. This entry into meiosis is termed the “transition zone” and the germ cells begin differentiating to form mature oocytes. The transition zone is classically identified by crescent-shaped DAPI morphology due to the polarized active movement of chromosomes; however, in certain mutant situations that affect chromosome pairing or germ cell proliferation, this region with distinct DAPI morphology may be either absent or extended (*e.g. hal-2* and *syp-1)* [4,5]. Following the transition zone, germ cell nuclei enter pachytene stage where chromosomes no longer are undergoing rapid polarized movement and instead assume a cage-like appearance. After pachytene, chromosome begin the condensation process in the diplotene stage and eventually fully condense to form six DAPI-staining bodies (one for each set of homologs) at diakinesis. This “pipeline” of germ cell development in the *C. elegans* gonad has enabled the visualization of all stages of germ cell development simultaneously within a single germline, thereby making this model system a powerful tool for cytological approaches.

Cytological studies of the *C. elegans* germline illuminate key aspects of meiosis, including chromosome pairing, recombination, regulation of DNA damage responses, and apoptosis in gamete production [6–8]. The spatial-temporal organization of the germline can be used to define the timing and/or progression of these events throughout meiotic prophase I in *C. elegans* [7–9]. For example, localization and quantification of foci composed of meiotic recombination proteins established the timing and steps of DNA repair events in the *C. elegans* germline [6,10–12]. Further, quantification of these foci within the germ cell nuclei can indicate changes in the frequency of these specific DNA repair events both in wild type and mutant contexts. Overall, quantitative image analysis of whole germlines have been instrumental in revealing roles for specific genes in meiotic DNA repair [8].

Germ cell differentiation and fertility in *C. elegans* require the germline to assemble RNA/protein condensates called P granules. These membraneless organelles are perinuclear during the majority of germ cell development and are involved in silencing germline transcription via small RNA pathways [13–15]. For nearly 40 years, cytology and genetics have played critical roles in studies of P granules. In 1982, P granules were originally identified by immunofluorescence imaging that revealed the existence of granules in the *C. elegans* P cell lineage, which exclusively gives rise to the germline [16]. Subsequent high-resolution microscopy, live imaging, and fluorescence recovery after photobleaching studies have revealed the components, dynamics, and liquid-like properties of P granules [13–15]. Further, analysis of whole adult gonads stained for P granule structures reveal that some components of these membraneless organelles can undergo morphological changes during meiotic prophase I progression [17], further suggesting possible changes in function during oogenesis.

While both qualitative and quantitative microscopy approaches are currently employed to study the *C. elegans* germline, the variation in the chromosome morphology throughout the germline and technical variability from affixing dissected gonads affixed to microscope slides have limited high-throughput automated analysis of germline features. Due to a lack of automated image analysis, many research groups rely on time consuming and laborious manual efforts for quantifying features of germ cells within whole *C. elegans* germlines. To expedite and expand quantitative image analysis of the entire *C. elegans* germline, we developed a high-content, automated method using custom algorithms that function with image processing software. This method enables quantitative image analysis of cytological features of single nuclei within whole *C. elegans* gonads. Further, this computational pipeline permits analysis and data visualization of individual nuclei based on their position within the germline. Here we describe and validate our computational method by analyzing images of multiple features of germ cell nuclei undergoing meiotic prophase I progression within the context of an entire *C. elegans* germline.

## Results

### Gonad Analysis Pipeline for fluorescent image analysis of whole *C. elegans* germlines

The *C. elegans* germline presents many challenges for automated quantification of cytological data. Due to the non-linear three-dimensional (3D) shape of both undissected and dissected gonads, it has been difficult to computationally: 1) distinguish individual nuclei within an imaged gonad; and, 2) contextualize quantitative features of individual nuclei based on their position in the gonad and during specific stages of meiotic prophase I. Further, the freedom of dissected gonads to adopt multiple shape confirmations when affixed to a microscope slide or coverslip, presents an additional challenge for automating computational analysis of large numbers of dissected gonads. To overcome these challenges, we constructed a Gonad Analysis Pipeline using image quantification software in conjunction with custom scripts implemented in MATLAB and R to enable high-throughput quantification of germline features at single nucleus resolution, while maintaining information regarding the relative position of these nuclei within the *C. elegans* germline. For all of our analyses, we acquired and utilized 3D immunofluorescence images of dissected, fixed *C. elegans* germlines using established protocols that preserve the 3D architecture of the germline (Figure 1A; Materials and Methods). Since high-resolution analysis of whole *C. elegans* gonads requires acquisition of multiple 3D images to encompass their entire distal-proximal length, we stitched the individual 3D images together into a single reconstruction of the imaged germline using either Imaris Stitcher or an image stitching plugin in FIJI (see Materials and Methods; [18]). Individual nuclei within the gonad were defined using Surface in Imaris with the DNA stain DAPI (see below). Due to the arrangement of nuclei in some germlines, a subset of nuclei (23%) were unable to be computationally identified and were subsequently removed from the dataset (Figure 1B). A caveat of removing these nuclei is that specific germline regions could be under sampled (p<0.001 Chi Square Test of Goodness of Fit, Supplemental Figure 1A); however, we found that combining the datasets of multiple germlines enabled even sampling of nuclei across the germline from the pre-meiotic tip to the end of late pachytene (p=0.422, Chi Square Test of Goodness of Fit, Supplemental Figure 1B). From our imaged gonads (which capture the top 25-30% of the germline along the dorsal-ventral axis; see Materials and Methods), we computationally identified an average of 146.3±16.9 nuclei per germline (n=4 gonads). Overall, these results indicate the ability of this pipeline to identify and analyze large numbers of nuclei from whole gonads.

**Figure 1.**
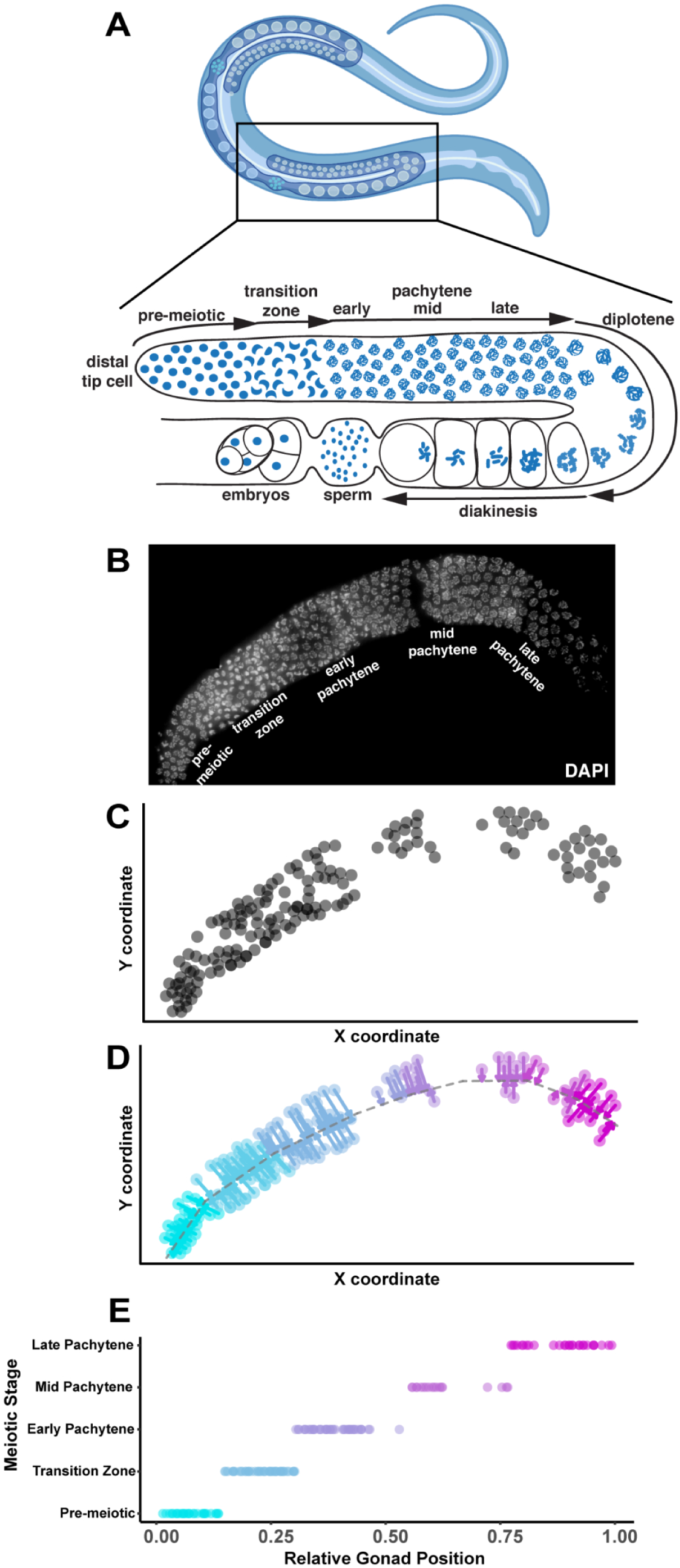
Gonad Linearization algorithm transforms and orients 3D-nuclei within a non-linear *C. elegans* gonad onto a one-dimensional axis. We designed a custom algorithm (called the “Gonad Linearization algorithm”) to enable the assessment of individual *C. elegans* nuclei relative to their position within a germline. (A) Cartoon of adult hermaphrodite worm (above panel; made with Biorender) with zoom in of one gonad arm (lower panel) with nuclei (blue) and indicated stages of meiosis based on DNA morphology (B) Dissected *C. elegans* hermaphrodite germline with DNA stained using DAPI (white). (C) 2D coordinate positions (units arbitrary) of individual whole nuclei (gray circles) within a *C. elegans* germline. Whole nuclei and respective coordinate positions were defined using Imaris. Nuclei found to be overlapping or only partially imaged were eliminated from analysis. Nuclei that were not able to be computationally oriented were also removed from analysis. (D) Application of the Gonad Linearization algorithm transforms the coordinates of nuclei onto a central axis line drawn through the germline, approximating the progression of nuclei through the germline based on their position along that line. (E) Normalizing the total length of line segments drawn through the center of the gonad enables standardized assessment of individual nuclei contextualized by their progression through the germline. Line segments were specifically placed to delineate the premeiotic zone and transition zone based on DAPI morphology of chromosomes. Early, mid, and late pachytene were defined on this graph by dividing the remaining normalized germline length into equal thirds.

To demarcate the conformation of each gonad from the distal tip (premeiotic) to proximal end (late pachytene), we drew contiguous line segments down the center of each germline (Figure 1C). This method allowed us to designate the different stages of meiotic prophase I along this segmented line based on DNA morphology: the premeiotic zone, transition zone (encompassing leptotene and zygotene), and pachytene. Since some mutant germlines lack some of these cytological features (*e.g.* absence of polarized chromosomes characteristic of transition zone nuclei), we developed an algorithm to approximate the relative germline position of each nucleus independent of DNA morphology (Figure 1D). This algorithm (called the “Gonad Linearization Algorithm”) approximates the position of each nucleus along the length of the germline based on its orientation relative to the line drawn along the center of the gonad. To calculate the position of each nucleus, the Gonad Linearization Algorithm identifies the best fit perpendicular intersection point for the position of each nucleus relative to the central line segments (see perpendicular arrows projecting from each nucleus to the central line in Figure 1C). This analysis allows us to recontextualize individual nuclei from 3D space into a one-dimensional (1D) space, enabling assessment of nucleus features based on position in the gonad as nuclei progress through meiotic prophase I.

To assess the ability of the Gonad Linearization Algorithm to accurately align nuclei through the germline, we applied the algorithm to a simulated dataset of 100 ‘germlines.’ Each simulated ‘germline’ contained 100 simulated ‘nuclei’ dispersed along the lengths of the ‘germline’ (Supplemental Figure 2). We found that, for most simulated ‘germlines,’ >90% of the ‘nuclei’ were accurately assigned to the correct line segment, and that correctly aligned nuclei recapitulated the order in which they were simulated along the length of the ‘germline’ (p<0.001, R2=1, Linear regression analysis, Supplemental Figures 2B,2C). Even in the case of incorrect assignment of a ‘nucleus’ to a line segment, the deviation of the placement of each ‘nucleus’ in the context of the whole gonad was <10% (Supplemental Figure 2D). In addition, we have included within the algorithm a way to manually correct the assignment of these incorrectly assigned nuclei. These data illustrate the accuracy and customizability of the Gonad Linearization Algorithm for analysis of diverse confirmations of dissected gonads.

### Quantification of DNA-associated proteins at single nucleus resolution

Manual quantification of foci within nuclei from whole *C. elegans* gonads is a laborious, rate-limiting step during image analysis. To validate our Gonad Analysis Pipeline’s automated quantification of meiotic features, we first quantified classic markers that are involved in double stand DNA break (DSB) formation and repair. The recombinase RAD-51 loads at sites of DSBs in meiotic nuclei [19,20]. The number of RAD-51 foci within germline nuclei can indicate either the extent of DSB induction and/or the efficiency of DSB repair during meiotic prophase I progression [21,22]. DSB-2 promotes DSB induction, and accumulates on meiotic chromatin in the final stages of the transition zone and early pachytene when RAD-51 forms numerous foci [22,23].

To quantify RAD-51 within an entire germline, we implemented our Gonad Analysis Pipeline adapted with a custom MATLAB script in combination with the Gonad Linearization Algorithm. First, we identified nuclei within the germline using DAPI (see Methods for details). A custom MATLAB script (called Spots to Surfaces) was used to: 1) identify the RAD-51 foci (spots) that were associated with each individual nucleus (surface); and 2) provide a readout of foci per nucleus. Then after drawing line segments along the length of the gonad, the Gonad Linearization Algorithm was used to transform the position each nucleus and the RAD-51 foci (spots) associated with that nucleus on to that 1D line. This transformation generated data from a single germline that contained both the number of spots associated with each nucleus and the relative position of each nucleus along the length of the germline. In addition to scoring the number of RAD-51 foci for each nucleus, we further calculated the mean intensity of DSB-2 staining with each nucleus using Imaris (Figure 2A). From these analyzes, we are able to observe the complete dynamics of DNA repair at a single nucleus resolution.

**Figure 2.**
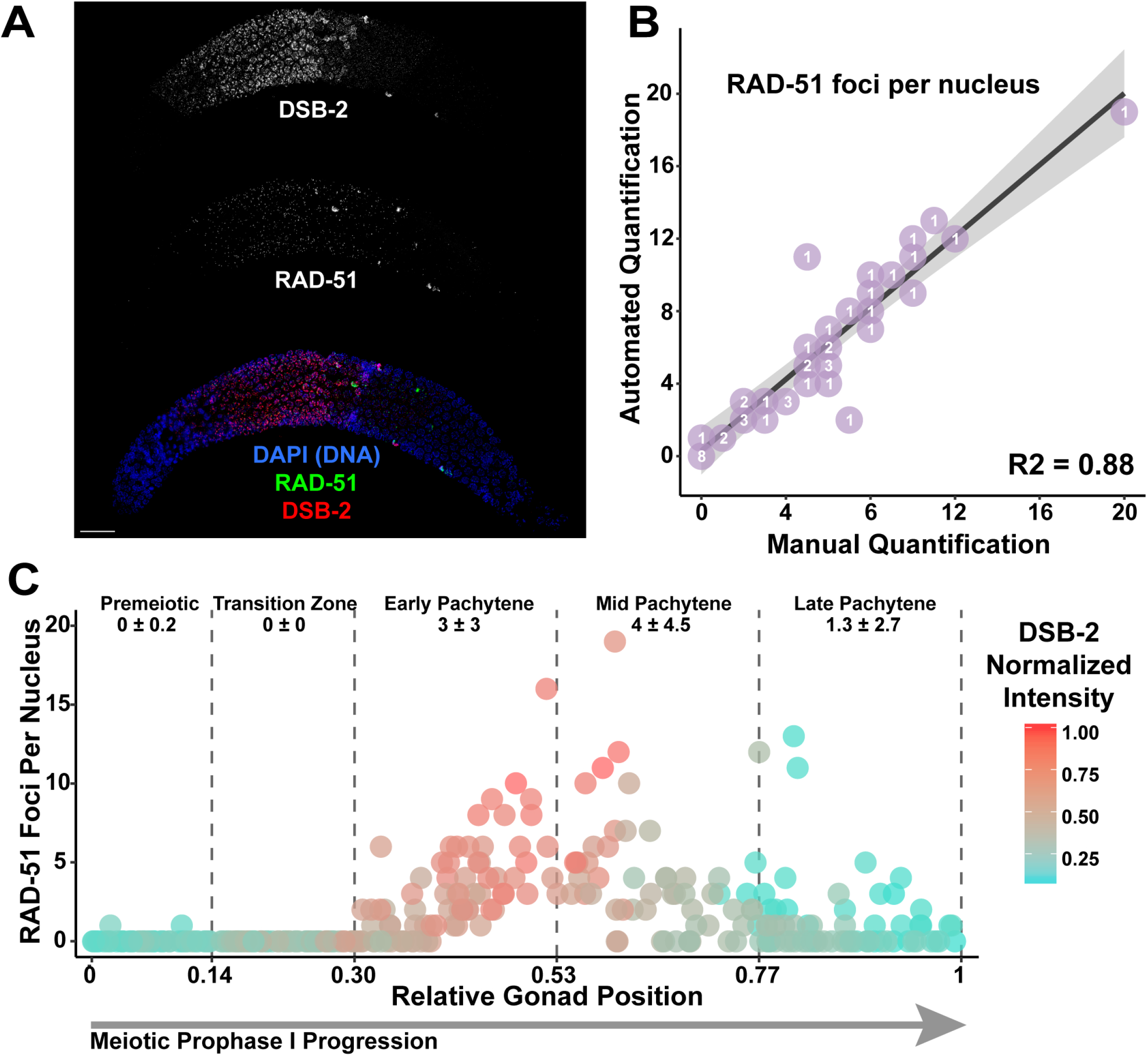
Computational pipeline enables germline-wide single nucleus assessment of double-strand DNA break (DSB) levels. (A) Immunofluorescence image of a *C. elegans* hermaphrodite germline stained with DAPI (DNA; blue), DSB-2 (red), and RAD-51 (green). Scale bar represents 20μm. (B) Comparison of data from automated quantification of RAD-51 foci associated with individual nuclei to data from manual quantification of RAD-51 foci within those same nuclei analyzed by the automated system (n=47). The number listed on each point (purple) indicates the number of nuclei scored with that result. The linear regression line is displayed as a black line, while the grey shaded area represents the 95% confidence interval of this analysis. (C) Visualization of RAD-51 foci counts and DSB-2 normalized mean fluorescence intensity of individual nuclei across gonads (n=2). The RAD-51 foci counts and DSB-2 normalized intensity values of 295 individual nuclei are displayed. DSB-2 mean intensity was normalized within analyzed gonads by the highest recorded DSB-2 mean fluorescence intensity among nuclei. Normalized DSB-2 intensity is indicated with a color gradient from red (highest intensity) to blue (lowest intensity). Vertical dashed lines indicate the average position in which nuclei within each gonad transition between each successive stage of meiotic prophase I (as indicated by text in the figure), as indicated by DAPI morphology. The deviation between these transition points was <0.01 between the germlines. Numbers below the text demarcating each respective meiotic stage in the germline indicate the mean number of RAD-51 foci ± the standard deviation of RAD-51 foci among nuclei within that region.

To determine the accuracy of our high-throughput Gonad Analysis Pipeline method for nuclear-associated foci quantification, we manually scored RAD-51 foci in a blinded subset of representative nuclei taken from whole gonad images (Figure 2B; n=47 nuclei). The mean deviation between automated and manual foci quantification was 0.06±1.45, and the number of foci per nucleus quantified by Imaris software correlated well with the number of foci scored manually (p<0.001, Adjusted R^2^ = 0.88, Linear Regression Analysis, Figure 2B). Thus, across a population of nuclei, our Gonad Analysis Pipeline yields reliable statistics for the number of foci associated with individual *C. elegans* germline nuclei.

Using the Gonad Analysis Pipeline, we assessed the relationship between DSB-2 and RAD-51 along the length of the germline (Figure 2C). In concordance with previous studies [19,22,23], we observe most nuclei with one or more RAD-51 foci within the central ~50% of the germline (Figure 2C), corresponding to the end of the transition zone through mid-pachytene stages of meiosis I (Figure 1D). The per-nucleus normalized mean intensity of DSB-2 within germlines was also highest in the central 50% of the germline (Figure 2C). To dissect this relationship further, we binned the DSB-2 and RAD-51 data into two bins based on when DSB-2 is loaded to chromatin in early prophase (transition zone-early pachytene) or offloaded from chromatin in late prophase (mid-late pachytene) [22]. Overall, higher DSB-2 intensity is correlated with increased numbers of RAD-51 foci (Supplemental Figure 3). Notably, we observed a stronger correlation in early prophase (Spearman’s *ρ* 0.785 95% CI 0.721-0.836, p value < 0.001, Spearman’s rank correlation test) than in late prophase (Spearman’s *ρ* 0.389 95% CI 0.225-0.532, p value < 0.001, Spearman’s rank correlation test), supporting the reported function of DSB-2 to promote DSB induction [22]. Further, this result demonstrates the capability of the Gonad Analysis Pipeline to quantify the relationships of cytological features at single nucleus resolution.

### Quantification of meiotic chromosome structure-associated foci at single nucleus resolution

Next we used the Gonad Analysis Pipeline to quantify foci associated with specific steps in DSB repair that occur along meiotic chromosome axis structures. While many proteins are involved in establishing a crossover during meiosis, we focused on quantifying the localization pattern of two proteins that are loaded after the initial strand invasion steps of recombination. The MutS homolog MSH-4/5 and cyclin-like COSA-1 localize to intermediate steps in the meiotic DSB repair process and are required for crossover recombination events between homologous chromosomes [11,19,24,25]. In early-mid pachytene, MSH-5 has been observed to form many dim foci before late pachytene, when both COSA-1 and MSH-5 localize to 6 foci, marking the positions of the obligate crossover for each of the six *C. elegans* chromosomes. Studies have demonstrated that the synaptonemal complex – a proteinaceous structure that assembles between homologous chromosomes during meiosis – recruits MSH-5 and COSA-1 in *C. elegans* [19,26–29,31].

We adapted Gonad Analysis Pipeline to determine the number of MSH-5 and COSA-1 foci associated with the synaptonemal complex protein, SYP-1 throughout the germline (Figures 3A, 3B). For this approach, SYP-1 staining was used instead of DAPI to generate surfaces for each individual nucleus. Next, we identified MSH-5 or GFP::COSA-1 foci, then used the Spots to Surface MATLAB script to identify the foci associated with each SYP-1 surface, and finally approximated the positions of these SYP-1 surfaces along the germline using the Gonad Linearization Algorithm. As the synaptonemal complex is not fully assembled until the end of the transition zone, we did not identify any SYP-1 objects in the first segmented portion of each analyzed germline, which corresponds to the pre-meiotic region (Figure 3C). In total, we identified the SYP-1 surfaces of 167 individual nuclei in a single germline stained with SYP-1 and MSH-5, and 168 individual nuclei in a single germline stained with SYP-1 and GFP::COSA-1. As previously reported [10,11], MSH-5 forms >6 foci per meiotic nucleus in early-mid pachytene. Then in late pachytene (the final ~25% of the germline), GFP::COSA-1 forms bright, robust foci and both MSH-5 and COSA-1 foci counts converge to ~6 foci per nucleus, which corresponds to the 6 total crossovers formed per nucleus [11]. This result demonstrates the capability of our approach to not only identify nuclear structures, but to quantitate the subnuclear association of specific meiotic proteins with specific chromosome structures at single-nucleus resolution.

**Figure 3.**
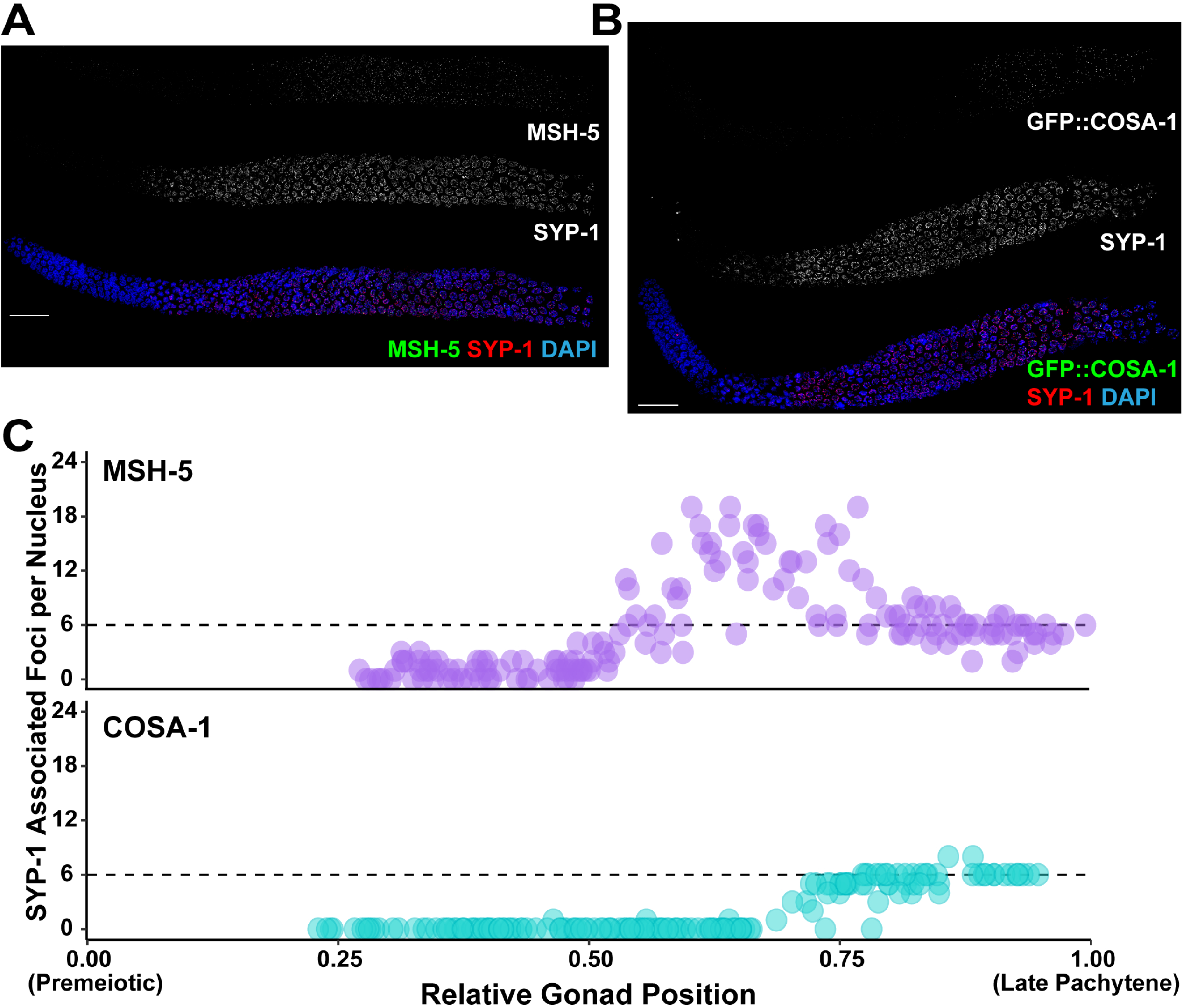
Single nucleus analysis of meiotic recombination markers along the meiotic chromosome axis. Immunofluorescence images of a *C. elegans* hermaphrodite germline stained with either (A) DAPI (DNA; blue), MSH-5 (green), and SYP-1 (red), or (B) DAPI (blue), GFP::COSA-1 (green), and SYP-1 (red). Scale bar represents 20μm. (C) Visualization of numbers of MSH-5 (purple) or GFP::COSA-1 (blue) foci associated with SYP-1 within individual nuclei across the germlines displayed in A-B. As nuclei progress through meiotic prophase I, the number of MSH-5 and COSA-1 spots converge at 6 foci per nucleus in the latter part of the germline, consistent with the reported number of MSH-5 and COSA-1 foci marking the 6 crossover sites in late pachytene [11].

### Quantification of perinuclear structures across the *C. elegans* germline

To demonstrate the ability of our method to assess extranuclear features of the *C. elegans* germline, we adapted our Gonad Analysis Pipeline to identify and quantify P granule structures that assemble within the perinuclear space of germ cells. P granules are liquid-like condensates associated with nuclear pore complexes in the *C. elegans* germline that process small RNAs [13]. For our analysis of P granules, we analyzed two components of P granules: PGL-1 and ZNFX-1. PGL-1 is a core component of P granules that is required for fecundity [16,30]. ZNFX-1 is a P granule component required for effective transcript silencing in the germline and colocalizes with PGL-1 perinuclear foci in the germline [32,33].

To analyze the localization of PGL-1 and ZNFX-1 P granule components throughout adult germline (Figure 4A), we adapted our Gonad Analysis Pipeline to initially identify and quantify the number of individual perinuclear PGL-1 and ZNFX-1 foci by creating surfaces of each focus in Imaris (Figure 4A). In total, we identified n=4779 PGL-1 foci and n=4034 ZNFX-1 foci (Figure 4B). Then, we applied the Gonad Linearization Algorithm to approximate the position of these foci relative to their progression through the germline (Figure 4B). To understand the relationship between PGL-1 and ZNFX-1, we determined the proportion of colocalized PGL-1 and ZNFX-1 along the in a sliding window representing 10% of total gonad length (Figure 4C). Throughout meiotic prophase I, >50% of PGL-1 and ZNFX-1 foci are consistently colocalized; however, in late prophase I, the frequency of colocalization increases to ~75%. From our analysis, we also found that PGL-1 foci were more frequently found unassociated with ZNFX-1 than ZNFX-1 was found unassociated with PGL-1 (Figure 4D). Together, these results agree with previous results indicating the colocalization of these two components within the *C. elegans* hermaphrodite germline [32,33]. Overall, this data demonstrates the adaptability and customizability of the Gonad Analysis Pipeline to quantitate the changes in colocalization frequency throughout the *C. elegans* germline.

**Figure 4.**
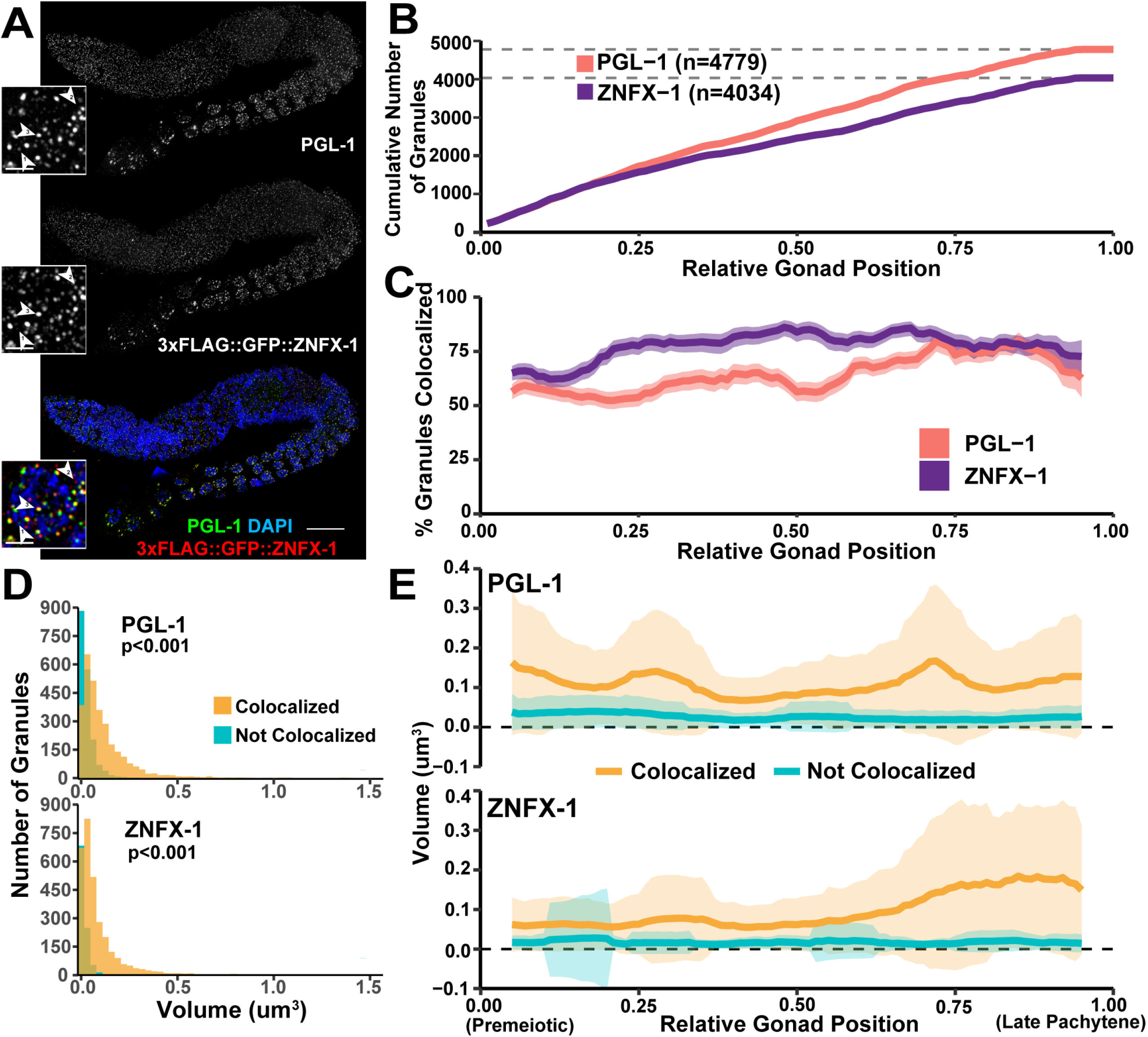
Assessment of P-granule components across meiotic prophase I. (A) Immunofluorescence image of a *C. elegans* hermaphrodite germline stained with DAPI (DNA), PGL-1 (green), and 3xFLAG::GFP::ZNFX-1 (red). Inset images show a single pachytene nucleus. Numbered arrowheads respectively indicate examples of: (1) a PGL-1 focus not colocalized with ZNFX-1, (2) a ZNFX-1 focus not colocalized with PGL-1, and (3) colocalized PGL-1 and ZNFX-1 foci. The scale bar in the whole germline image represents 20μm, while the scale bars in the insets represent 2μm. (B) Cumulative number of PGL-1 and ZNFX-1 foci identified across the germline. (C) Percent of total PGL-1 and ZNFX-1 foci which are respectively colocalized within a sliding window representing 10% of total germline length. Shaded area represents 95% Binomial Confidence Interval. (D) Histograms displaying the distribution of PGL-1 and ZNFX-1 foci volumes, distinguishing between foci colocalized (yellow) or not colocalized (blue) with other respective protein. P values were calculated from comparisons between colocalized and non-colocalized focus volumes by Mann-Whitney U test. (E) Mean volume of PGL-1 and ZNFX-1 foci in a sliding window representing 10% of total germline length, distinguishing between foci which are (yellow) or are not (blue) colocalized. Shaded area represents standard deviation.

To test whether our method could quantify additional structural features of P granules, we quantified and compared the volume/size of individual PGL-1 and ZNFX-1 P granules to the volume/size of P granules with colocalized PGL-1 and ZNFX-1. From our analysis across meiotic prophase I, we found that the volume of foci that were colocalized were larger than individualized foci for both proteins assessed (p<0.001, Mann-Whitney U test, Figure 4D). When we examined the mean volume of PGL-1 and ZNFX-1 foci in a sliding window representing 10% of total gonad length (Figure 4E), we observed that P granules with colocalization of PGL-1 and ZNFX-1 were consistently larger in volume than those granules that did not have both components present. This result may indicate that the inclusion of multiple P granule components possibly results in a synergistic increase the volume of a granule. Taken together, we have demonstrated that our approach enables high-throughput analysis of germline granules provides support for a model in which the composition and features of individual P granules may change throughout meiotic prophase I progression.

## Discussion

In this study, we demonstrate the utility of a customizable computational pipeline, called the Gonad Analysis Pipeline, developed to perform automated quantification of features within (or associated with) individual nuclei with reference to the position of the nuclei in the *C. elegans* gonad. Specifically, we adapt and use Gonad Analysis Pipeline to quantify foci per nucleus, foci associated with chromosome structures, and foci colocalization frequencies across whole adult *C. elegans* hermaphrodite gonads from the pre-meiotic tip to late pachytene. This pipeline yields datasets concordant with previous observations for known features of meiotic prophase I. Additionally, many *C. elegans* mutants defective in key meiotic events such as synapsis and pairing can have aberrant DNA morphology and disruption of normal meiotic stage progression. These defects make it difficult to use DNA morphology to discern the specific transitions between meiotic stages and challenging to categorically delineate nuclei within those germline contexts. Our automated Gonad Analysis Pipeline provides a consistent metric utilizing position along the normalized gonad length for comparative analysis of mutants to wildtype germlines.

While analyses presented here assess nuclei from the pre-meiotic tip to late pachytene of the *C. elegans* germline, our pipeline can also be extended to include more proximal portions of the germline for quantitative analyses of other germline features. For example, P granules display a dynamic localization pattern throughout the germline, changing from cytoplasmic localization in the distal region of the germline to a more perinuclear localization in the more proximal region of germline [13]. Our computational pipeline can be utilized to quantify these changes in P granule localization across the entire *C. elegans* germline and perform comparative studies of these nucleus-cytoplasm localization dynamics between wild type and mutant contexts. Additionally, several studies have found dynamic changes to the localization of specific synaptonemal complex components during meiotic prophase progression [34–37]. Our pipeline can also be utilized to quantify these changes in the chromosome axis and the synaptonemal complex from transition zone through diakinesis.

Our analyses demonstrate how small customizable changes to the Gonad Analysis Pipeline can enable quantification at multiple levels from the entire germline to single nuclei. Additional changes can enable the additional quantifications of cytological objects, such as sphericity, intensity, and relative distance between objects. Utilization of these other quantifiable metrics enable a comprehensive analysis of many germ cell features, including the quantification of chromosome pairing for fluorescence in situ hybridization (FISH) experiments, assembly and disassembly of chromosome structures, and protein dynamics during live cell imaging. In particular for live imaging, the pipeline could assess changes in numerous metrics such as velocity, mean square displacement, duration, volume, and sphericity of objects over time for all nuclei during oogenesis and contextualize these statistics based on nuclear position within the germline. These types of adaptations of our Gonad Analysis Pipeline for live imaging may prove particularly powerful for quantification of the liquid-like properties and dynamics of P granules in the adult germline, especially in response to different stresses or aging.

The present study focuses on adult hermaphrodite germlines, however, the Gonad Analysis Pipeline can also be used to analyze larval germlines and adult male germlines. An increasing number of studies are demonstrating the power of performing comparative analyses between oogenesis and spermatogenesis in *C. elegans* to identify important sexual dimorphic features of meiosis [6,9,38,39]. Spermatogenesis in the germlines of *C. elegans* males is also organized in a spatial-temporal gradient [40] and can easily be analyzed by our pipeline, thereby aiding both studies of spermatogenesis as well as sexual dimorphism of germ cell development.

Taken together, we have generated and validated an automated and customizable image analysis resource for the *C. elegans* germline community. Our Gonad Analysis Pipeline enables standardized quantification of diverse features of the *C. elegans* gonad. Moreover, our approach is flexible and could be applied to analyze features of other tissues composed of cells organized along a linear gradient.

## Acknowledgements

We thank A. Villeneuve for antibodies, the CGC (funded by National Institutes of Health (NIH) P40 OD010440) for strains, and Adele Adamo for providing the pET30a plasmid. We thank the members of the Libuda Lab for many insightful and helpful discussions in refining this method. We also thank A. Villeneuve for encouraging us to turn this work into a manuscript. This work was supported by the National Institutes of Health T32GM007413 to ET and AD; National Institutes of Health F32GM130006 to NAK; Jane Coffin Childs Postdoctoral Fellowship to CKC; and National Institutes of Health R35GM128890 Award to DEL. DEL is also a recipient of a March of Dimes Basil O’Connor Starter Scholar award and Searle Scholar Award.

---

**Supplemental Figure 1.**
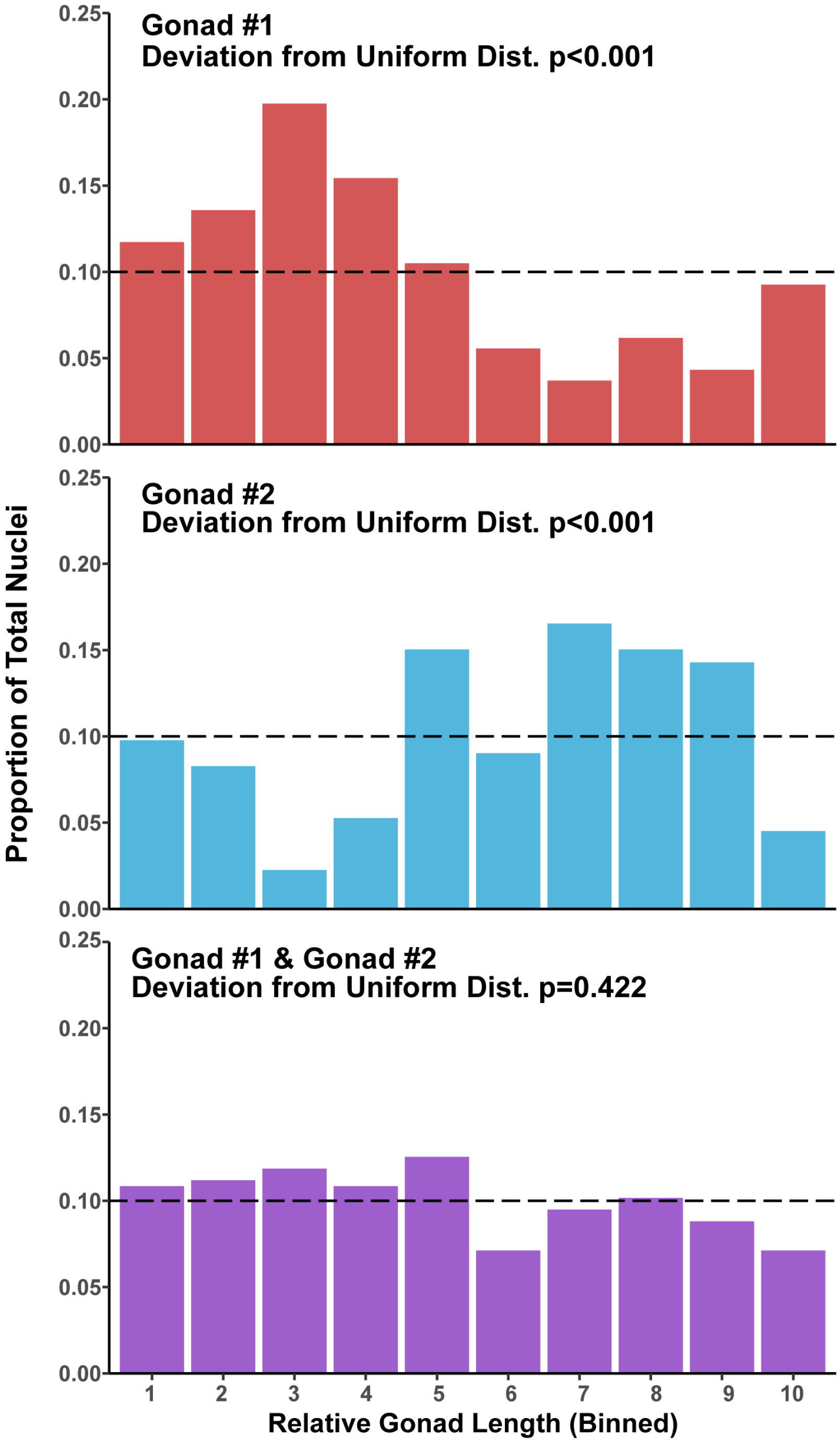
Gonad length distribution of whole nuclei identified as analyzable in individual germlines. Bar plots representing the proportion of nuclei in ten equal bins across the lengths of the two gonads analyzed in Figure 2. P values were calculated by Chi Square Test of Goodness of Fit (expected frequency 0.1 in each bin). The distribution of nuclei within bins is indistinguishable from a uniform distribution by this same test when the nuclei from the two germlines are taken together.

**Supplemental Figure 2.**
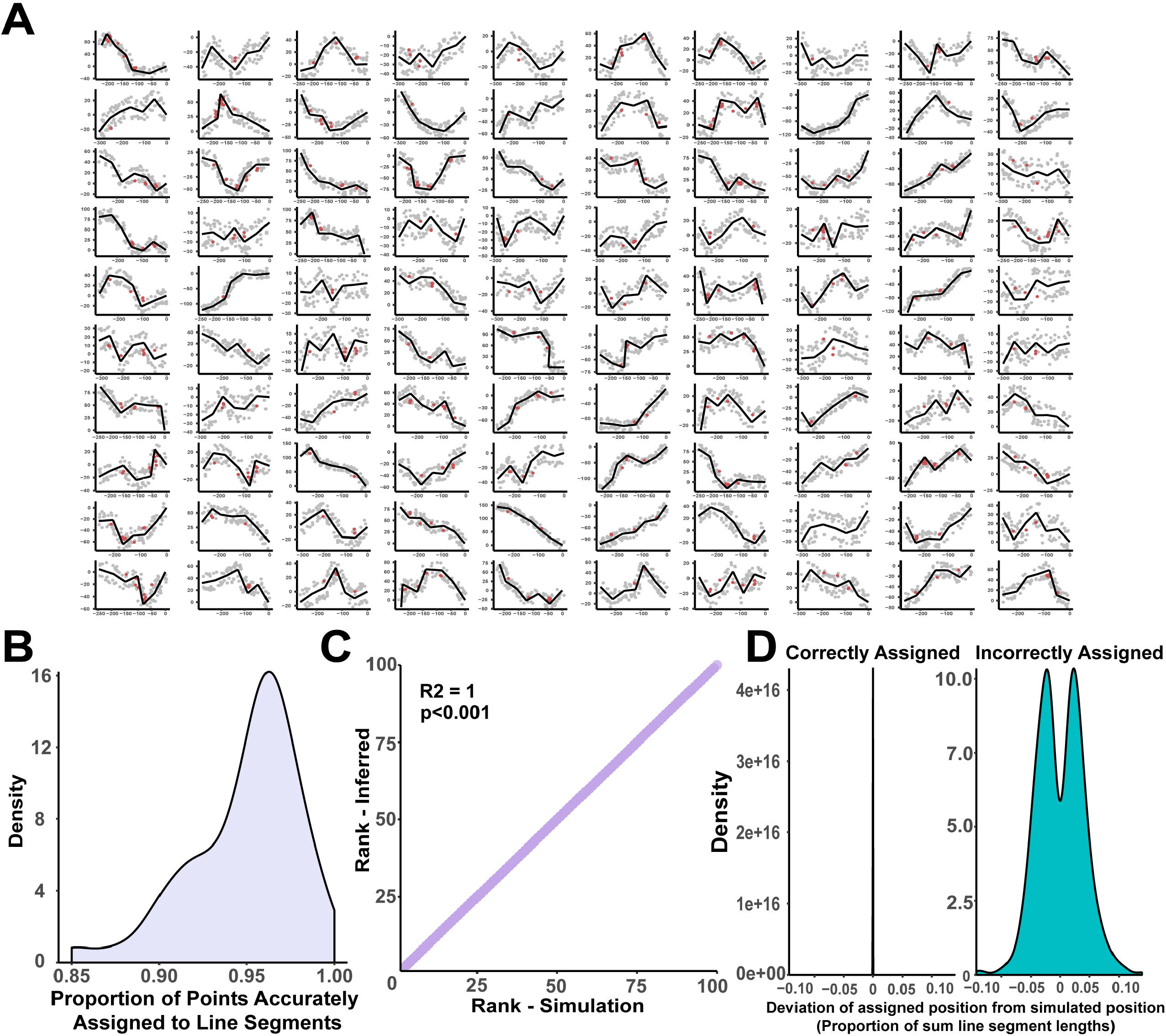
Validation of Gonad Linearization algorithm. (A) Plots of simulated dataset of 100 ‘germlines’ each with 100 ‘nuclei’ points dispersed along their lengths. Points were realigned to the central lines using the Gonad Linearization algorithm, and points that were aligned to the correct line segment are marked in grey while points marked in red were aligned to the incorrect line segment. (B) Density plot demonstrating the distribution of accuracy of point alignment to line segments among the 100 individual simulated ‘gonads’. (C) Comparison of the known rank order of correctly aligned spots to the rank order of spots as determined by the Gonad Linearization algorithm. R2 and p values were calculated by linear regression analysis. (D) Calculation of the deviation of assigned positions as determined by the Gonad Linearization algorithm from ‘actual’ known positions from the original simulation.

**Supplemental Figure 3.**
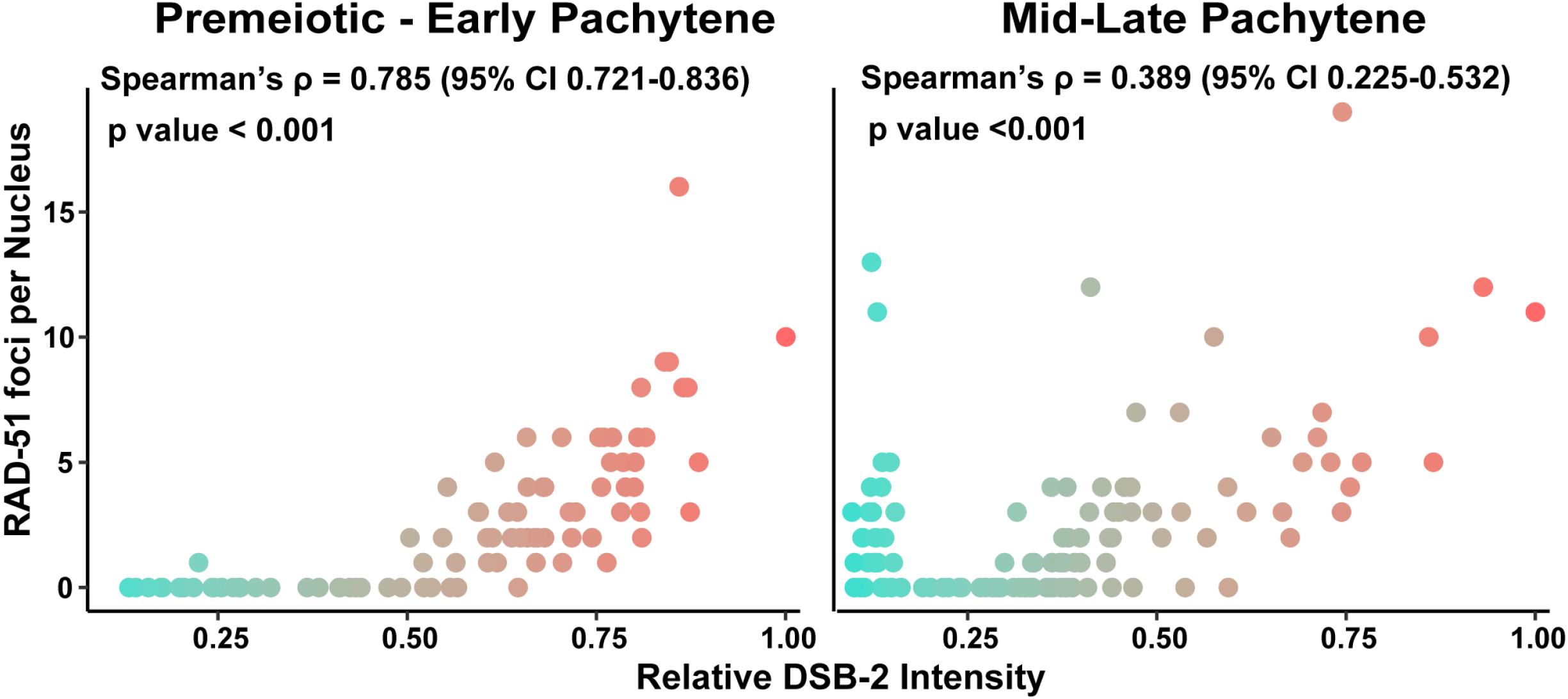
DSB-2 normalized intensity per nucleus correlates with RAD-51 foci per nucleus. Assessment of nonparametric correlation by Spearman correlation tests between RAD-51 foci per nucleus and normalized DSB-2 staining intensity among nuclei within the premeiotic through early pachytene stages, and in mid-through late pachytene stages.

## Materials and Methods

### *Caenorhabditis elegans* strains and maintenance

*C. elegans* strains were maintained under standard conditions on nematode growth medium (NGM) plates at 20°C with OP50 *Escherichia coli* bacteria lawns. All experiments were performed in the N2 background of *C. elegans* (CGC).

Strains used in this study include AV630 (meIs8[*unc-119(+) pie-1p*::GFP::*cosa-1*] II), N2 (wild type), and YY916 (*znfx-1(gg544[3xflag::GFP::znfx-1])* II.).

### Immunofluorescence

Immunofluorescence was performed as in Libuda *et al.* 2013 [27]. At 18-22 hours before dissection, L4 stage hermaphrodite worms were isolated and maintained at 20°C on NGM plates seeded with OP50. Gonads were dissected in 30μL of egg buffer (118mM NaCl, 48mM KCl_2_, 2mM CaCl_2_, 2mM MgCl_2_, 25mM HEPES pH7.4, 0.1% Tween20) and were fixed in egg buffer with 1% paraformaldehyde for 5 minutes on a Superfrost Plus slide (VWR). Gonads were then flash frozen in liquid N_2_ and the cover slip was removed. For germlines stained for DSB-2, RAD-51, MSH-5, or GFP::COSA-1, the slide was placed in −20°C MeOH for 1 minute and then was washed in PBST (1x PBS, 0.1% Tween20). For germlines stained for PGL-1 and ZNFX-1, the slide was placed in the slide was placed in −20°C MeOH for 10 minutes, then in −20°C acetone for 5 minutes, and then was washed in PBST. Slides were washed 3x in 1xPBST for 5 minutes before being place in block (1xPBS, 0.1% Tween20, 0.7% Bovine Serum Albumin) for 1 hour. 50μL of diluted primary antibody (in 1xPBST; see below for individual antibody concentrations) was applied to each slide and allowed to stain overnight in a dark humidifying chamber with a parafilm coverslip. At 16-18 hours after application of primary antibody, slides were washed 3x in PBST for 10 minutes. 50μL of diluted secondary antibody (in 1xPBST; see below for individual antibody concentrations) was applied to each slide and allowed to stain for 2 hours in a dark humidifying chamber with a parafilm coverslip. Slides were washed 3x in PBST for 10 minutes in a dark chamber and then 50μL of 2μg/mL of DAPI in ddH_2_O was added to each slide and incubated for 5 minutes in a dark humidifying chamber with a parafilm coverslip. Slides were washed in PBST for 5 minutes in a dark chamber and then were mounted in VectaShield with a No. 1.5 coverslip (VWR) and sealed with nail polish. Slides were maintained at 4°C prior to imaging (as described below). The following primary antibodies were utilized in this study at the listed concentrations: polyclonal chicken αRAD-51 (1:1000, this study, see below), αDSB-2 (1:5000; [22]), αMSH-5 (1:10,000, Novus #3875.00.02), polyclonal chicken αGFP (1:2000, Abcam #ab13790), monoclonal mouse αPGL-1 K76 (1:20, Developmental Studies Hybridoma Bank), polyclonal guinea pig SYP-1 (1:250; [28]), and polyclonal rabbit GFP (1:1000; [11]). Secondary staining was performed with goat antibodies conjugated to Alexa Fluors 488 and 555 targeting the primary antibody species (1:200, Invitrogen).

### Antibody production

Our RAD-51 antibody was generated from a His-tagged fusion protein expressed by Genscript from plasmid pET30a containing the entire RAD-51S coding sequence (1385 bp, GenBank accession number AF061201 [19,41]). Antibodies were produced in chicken and affinity purified by Pocono Rabbit Farms.

### Image Acquisition

Immunofluorescence slides were imaged at 512 × 512 or 1024 × 1024 pixel dimensions on an Applied Precision DeltaVision microscope with a 63x lens and a 1.5x optivar. To ensure analysis of the highest resolution germline images, we imaged the top ~quarter of the germline along the dorsal-ventral axis that encompassed whole nuclei closest to the coverslip, but our pipeline can be utilized for analysis of gonads imaged through entire dorsal-ventral axis. Images were acquired as Z-stacks at 0.2 μm intervals and deconvolved with Applied Precision softWoRx deconvolution software.

### Gonad Analysis Pipeline

Below is a detailed section describing the method. For a step-by-step protocol, please go to the publication section of www.libudalab.org.

#### Identification of nuclei within whole gonad images

3D images were tiled using the Imaris Stitcher software (Bitplane) or the Grid/Collection Stitching plugin in FIJI with regression threshold of 0.7 (this value was raised or lowered depending on the stitching results) [18]. If images were not accurately aligned by the Imaris Stitcher algorithm, they were manually adjusted before proceeding with analysis. Individual nuclei within stitched gonads were identified by DAPI as Surface objects. When using DAPI staining to define Surface objects, the changing morphology of nuclei within the germline required different sets of parameters to be utilized. Nuclei spanning from the distal premeiotic tip through the final 5 rows of pachytene were defined using Smooth 0.15, Background 3.5, Seed Point Diameter 2-3, and Volume Filter 8-55. Late pachytene nuclei (nuclei in the 5 rows preceding diplotene) were defined using Smooth 0.15, Background 4, Seed Point Diameter 3-4, and Volume Filter 10-50. Manual thresholding and specific values for Seed Point Diameter and Volume Filter were defined for each gonad within the indicated ranges. Defined Surfaces were then split to designate individual nuclei using the Imaris Surfaces Split module. Nuclei which were either partially imaged or overlapping with another nucleus, were eliminated from analysis.

#### Identification of SYP-1 surfaces in whole gonad images

In 3D stitched gonad images (see ‘Identification of nuclei within whole gonad images’, above) Individual SYP surfaces were defined using Absolute Intensity (enabled), Smooth (0.22), Background (N/A), Seed Point Diameter (N/A), and Volume Filter (deleted surfaces less than 0.5um). If multiple individual surfaces were generated to represent the SYP-1 staining of a single given nucleus, then these surfaces were manually unified.

#### Quantification of DSB-2 normalized mean staining intensity

DSB-2 mean staining intensity per nucleus was calculated using Imaris following definition of single nuclei as surface objects using DAPI signal (see “Identification of nuclei within whole gonad images” section). As image acquisition settings differed between imaged germlines but were consistent within the same germline, the DSB-2 mean intensity of each nucleus was normalized by dividing the mean intensity of each nucleus by the highest mean intensity among nuclei within a gonad.

#### Quantification of meiotic recombination foci

RAD-51, MSH-5, and GFP::COSA-1 foci were defined from stitched whole gonad images (see “Identification of nuclei within whole gonad images” section) using the Create Spots tool in Imaris (Bitplane) with the settings Estimated XY Diameter 0.1, Model PSF-elongation 1.37, and Background Subtraction enabled. To determine the number of RAD-51 foci per nucleus by determining based on proximity of defined Spots to Surfaces, we used a custom “Finds Spots Close to Surface” MATLAB module (Threshold value 1; see “Data and Code Availability” section for link to download module). The number of SYP-1 associated MSH-5 or GFP::COSA-1 foci per nucleus was also determined using the “Finds Spots Close to Surface” module (Threshold value 0.1).

#### Quantification of PGL-1 and ZNFX-1 foci

PGL-1 and ZNFX-1 foci were defined as Surface objects in Imaris (Bitplane) with the settings Smooth (Not enabled), Background 0.513, Seed Point Diameter (Not enabled), and Volume Filter (foci>0.1uM). In late pachytene, the large variance in different P granule sizes required the generation of a separate additional set of “large” surfaces with the settings Smooth (Not enabled), Background 0.513, Seed Point Diameter (Not Enabled), and Volume Filter A (0.1um - 2um) Filter B (0.1um - 12um). To ensure that moderately sized PGL-1 and ZNFX-1 foci were not counted twice in this analysis, we used the Surface-Surface Colocalization Xtension to identify overlapping ‘small’ and ‘large’ PGL-1 and ZNFX-1 foci respectively and generated a new intensity with values unique to colocalization surfaces. If two granules were found to be colocalized (shared the same unique intensity value), the foci from the ‘large’ analysis was removed from the dataset and the ‘small’ granule was kept, as these smaller granules better represented the images. Colocalization between PGL-1 and ZNFX-1 surfaces was similarly determined using the Surface-Surface Colocalization Xtension in Imaris and unique colocalization identity intensity channels.

### Gonad Linearization algorithm

To assess nuclei based on their position within the gonad, we used an algorithm (called “Gonad Linearization” algorithm) implemented in R to approximate the progression of nuclei through the *C. elegans* germline as a linearly ordered sequence beginning at the premeiotic tip and terminating at the end of pachytene. For link to download the Gonad Linearization algorithm, see “Data and Code Availability” section of Methods. To delineate the orientation of the gonad, a series of connected line segments marking the approximate center of the gonad were drawn on the stitched germline image using the Imaris Measurement tool. Specific measurement points were placed at positions indicating transitions between meiotic stages from DAPI nuclei morphology, specifically marking the beginning of the premeiotic zone, transition zone, pachytene, and end of pachytene.

Each line segment drawn through the germline was defined by the coordinates of its respective start (*x*_*i*_,*y*_*i*_) and end (*x*_*j*_,*y*_*j*_) points. The standard equation [0=*Ax* + *By* + *C*] of each line segment 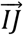 was calculated such that:

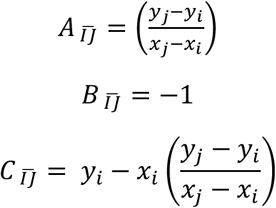

To determine whether the position of a nucleus within the gonad could be well approximated as a position on a given line segment, we calculated the perpendicular intersection point of a vector drawn from the position of the nucleus to each line segment. The perpendicular intersection point (*x*_*p*_,*y*_*p*_) of a nucleus at position (*x*_*n*_,*y*_*n*_) to a line 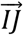 was calculated as:

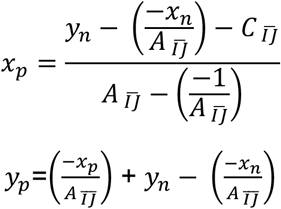

The transformed coordinate position (*x*_*p*_,*y*_*p*_) of a nucleus was considered well approximated if the distances from the start position of the line segment (*x*_*i*_,*y*_*i*_) to (*x*_*p*_,*y*_*p*_) and the distance from the end position of the line segment (*x*_*j*_,*y*_*j*_) to (*x*_*p*_,*y*_*p*_) were smaller than the total length of the line segment 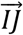. If multiple line segments met this criteria, the correct line segment was inferred to be the one for which the distance from the nucleus’ original position (*x*_*n*_,*y*_*n*_) to its perpendicular intersection point (*x*_*p*_,*y*_*p*_) was the shortest.

The above method of assigning nuclei to segments was sufficient for all germlines analyzed in this study. However, the specific arrangement of nuclei around the central gonad axis in the context of the whole germline conformation may lead to nuclei being incorrectly aligned according to these criteria. To ameliorate this potential problem, we included a stringency parameter in our algorithm, which increases the permissible distance nuclei may be assigned to a particular line segment. If increasing the stringency parameter from its default value of 0 is not sufficient to enable more accurate nuclei assignment, nuclei can also be manually assigned to line segments.

Once all nuclei had been assigned transformed coordinate positions, the sum length of all contiguous line segments drawn through a germline, as well as the sum distance of all line segments from the most proximal point to each transformed nucleus position, were calculated. Each length measurement was normalized to the total length of all line segments drawn through the germline to calculate relative gonad position, where a position of 0 corresponded to the start of the premeiotic tip and position 1 corresponded to the end of late pachytene.

### Validation of nucleus positioning by the Gonad Linearization algorithm

100 ‘gonads’ were simulated by iteratively generating six consecutive line segments with lengths ~Normal(50,5) and angles of intersection ~Normal(180,30). 100 points were simulated along the sum length of the line segments for each gonad ~Uniform(0,sum line segment lengths). Each point was then transposed perpendicularly to its line segment a distance ~Normal(10,3). These transposed ‘nucleus’ positions were then realigned to the line segments using the Gonad Linearization algorithm and were subsequently analyzed to determine goodness of fit.

### Statistics

All statistics were calculated in R (v3.5.1). Data wrangling was performed using the Tidyverse package (v1.3.0). Nonparametric correlations between DSB-2 normalized staining intensity and RAD-51 focus counts (Supplemental Figure 3) were assessed by Spearman correlation tests with confidence intervals calculated using the DescTools package (v0.99.30). Comparisons of RAD-51 focus manual and automated quantification (Figure 2B) and the rank order of simulated nucleus position data (Supplemental Figure 1C) were performed by linear regression analysis. The 95% Binomial confidence interval for the proportion of colocalized PGL-1 and ZNFX-1 granules (Figure 4C) was calculated using the DescTools package. Volumes of PGL-1 and ZNFX-1 (Figure 4D) foci were compared by Mann-Whitney U test.

### Data and Code Availability

All strains and antibodies available upon request. A step-by-step protocol for the Gonad Analysis Pipeline can be found at www.libudalab.org in the publication section. The “Gonad Linearization” algorithm and “Finds Spots Close to Surface” MATLAB module are available at github.com/libudalab/Gonad-Analysis-Pipeline. Figure S1 displays bar plots representing the proportion of nuclei identified from each region of the germline by the Whole Gonad Pipeline. Figure S2A displays plots of the simulated ‘germlines’ and ‘nuclei’ utilized to validate the Gonad Linearization algorithm. Figure S2B displays a density plot of the proportion of ‘nuclei’ in simulated ‘germlines’ which were accurately assigned to central line segments. Figure S2C displays a plot comparing the rank order of simulated ‘nuclei’ correctly assigned to central line segments within simulated ‘germlines’ to their known simulated rank order. Figure S2D displays density plots showing the relative deviation of simulated ‘nuclei’ from their known simulated positions relative to the alignment performed by the Gonad Linearization algorithm. Figure S3D displays dot plots assessing the association of DSB-2 staining intensity and RAD-51 focus counts in germlines analyzed by the Gonad Analysis Pipeline. Supplemental material (Figures S1, S2, and S3) are available at Figshare.

